# Zero-shot segmentation using embeddings from a protein language model identifies functional regions in the human proteome

**DOI:** 10.1101/2025.03.05.641584

**Authors:** Ami G. Sangster, Cameron Dufault, Haoning Qu, Denise Le, Julie D. Forman-Kay, Alan M. Moses

## Abstract

The biological function of a protein is often determined by its distinct functional units, such as folded domains and intrinsically disordered regions. Identifying and categorizing these protein segments from sequence has been a major focus in computational biology which has enabled the automatic annotation of folded protein domains. Here we show that embeddings from the unsupervised protein language model ProtT5 can be used to identify and categorize protein segments without relying on conserved patterns in primary amino acid sequence. We present Zero-shot Protein Segmentation (ZPS), where we use embeddings from ProtT5 to predict the boundaries of protein segments without training or fine-tuning any parameters. We find that ZPS boundary predictions for the human proteome are more consistent with reviewed annotations from UniProt than established bioinformatics tools and ProtT5 embeddings of ZPS segments can categorize folded domains, sub-domains, and intrinsically disordered regions. To explore ZPS predictions, we introduce a new way to visualize protein embeddings that closely resembles diagrams of distinct functional units in protein biology. Since ZPS and segment embeddings can be used without training or fine-tuning, the approach is not biased towards known annotations and can used to identify and categorize unannotated protein segments. We used the segment embeddings to identify unannotated mitochondrion targeting signals and SYGQ-rich prion-like domains, which are functional regions within intrinsically disordered regions. We expect the protein segment organization revealed here to lead to valuable information about protein function, including about intrinsically disordered regions and other less understood protein regions.

## Introduction

Proteins can contain a variety of distinct functional units including independently folding regions, known as folded domains, and regions lacking stable structure, such as intrinsically disordered regions (IDRs). Together, these distinct functional units determine the overall function of the protein. Powerful bioinformatics tools, such as Pfam [1] and Prosite [2], are widely used to identify and categorize folded domains from amino acid sequences alone. Other approaches are applied to protein structures to identify folded domains within proteins, referred to as protein segmentation [3,4]. In addition to folded domains, IDRs are present in over 60% of human proteins [5], perform important functions [6], and can be identified directly from amino acid sequences with high accuracy [7]. Unlike folded domains, functional categorization of IDR types remains challenging, though recent works have shown some success using conserved sequence properties [8,9] and supervised deep learning [10,11]. Unsupervised bioinformatics approaches, such as fLPS2 [12] and a Chi-Score Analysis [13], can segment proteins more generally based on statistical properties of amino acid sequences. These approaches can distinguish IDRs from domains and have revealed a non-random modular architecture within IDRs [13].

Here we explore the use of Protein Language Models (pLMs) to identify and categorize folded domains and IDRs from the human proteome. We use the pLM ProtT5 [14] to encode protein sequences into high-dimensional vectors known as embeddings. ProtT5 was pre-trained through unsupervised learning, a process where the model predicts masked (or hidden) amino acids in protein sequences and therefore does not require labelled data. Performing this task requires ProtT5 to develop contextualized embeddings of each amino acid based on the surrounding protein sequence, which are then used to predict the masked amino acids. After pre-training, visualizations of dimensionally reduced embeddings can reveal associations with amino acid properties, structure, and evolutionary relationships [14–16]. Additionally, pLMs can be used as a starting point for further training, known as fine-tuning, or the embeddings from a pre-trained model can be used as input to train another model. Both options have been applied to a variety of protein related tasks [15–20], including amino acid level domain annotation [21].

Embeddings or pLMs can also be used to make predictions on a new task, different from the pre-training task, without training or fine-tuning on the new task. This is referred to as “zero-shot” prediction. Protein embeddings have been used for zero-shot prediction of variant effects [22,23] and drug-target binding [24]. Zero-shot applications offer the potential for biological discovery without being tied to the original training task or being biased by the content and availability of training data. Recent work in image analysis has demonstrated that an unsupervised vision transformer can identify the boundaries of objects in images without human labelled examples; this is referred to as zero-shot image segmentation [25,26]. Inspired by this work, we sought to test whether a transformer-based pLM, ProtT5 [14], can perform zero-shot protein segmentation. Here, we demonstrate that a change point analysis applied to ProtT5 embeddings can identify the boundaries of biologically meaningful protein segments. We achieved this without training or fine-tuning any parameters, so we refer to this as Zero-shot Protein Segmentation (ZPS). We compare ZPS boundary predictions to annotations from UniProt [27], a protein annotation database which curates annotations from the literature and combines annotations from other databases and bioinformatics tools such as MobiDB [28] and ProRule [2]. We show that for the human proteome, the segments defined by ZPS match the boundaries of annotations from MobiDB for disorder and compositional biases more closely than other unsupervised methods designed specifically for these types of protein sequences [12,13]. Furthermore, we find that the ProtT5 embeddings of protein segments defined by ZPS can be used to categorize domains, sub-domains, and types of IDRs. Finally, as a specific application of these approaches, we identify unannotated mitochondrion targeting signals and SYGQ-rich prion-like domains in the human proteome.

## Results

### Known domains, IDRs, and motifs are found within ProtT5 embedding of FUS

To investigate whether ProtT5 embeddings can be used to identify the boundaries of biologically meaningful protein segments, we first visualized the ProtT5 embedding of human RNA-binding protein FUS [14] as a heatmap (Fig 1A). We selected FUS because it is a well-studied protein with a combination of annotated folded domains and IDRs. In the heatmap, we observed distinct segments in the embedding along the protein’s amino acid sequence. We identified boundaries of these segments using a change point analysis, which we refer to as Zero-shot Protein Segmentation (ZPS) (see Methods) and then visualized them (Fig 1B). The segments defined by ZPS closely matched with annotations from UniProt and the literature [29,30] (Fig 1C).

**Fig 1.**
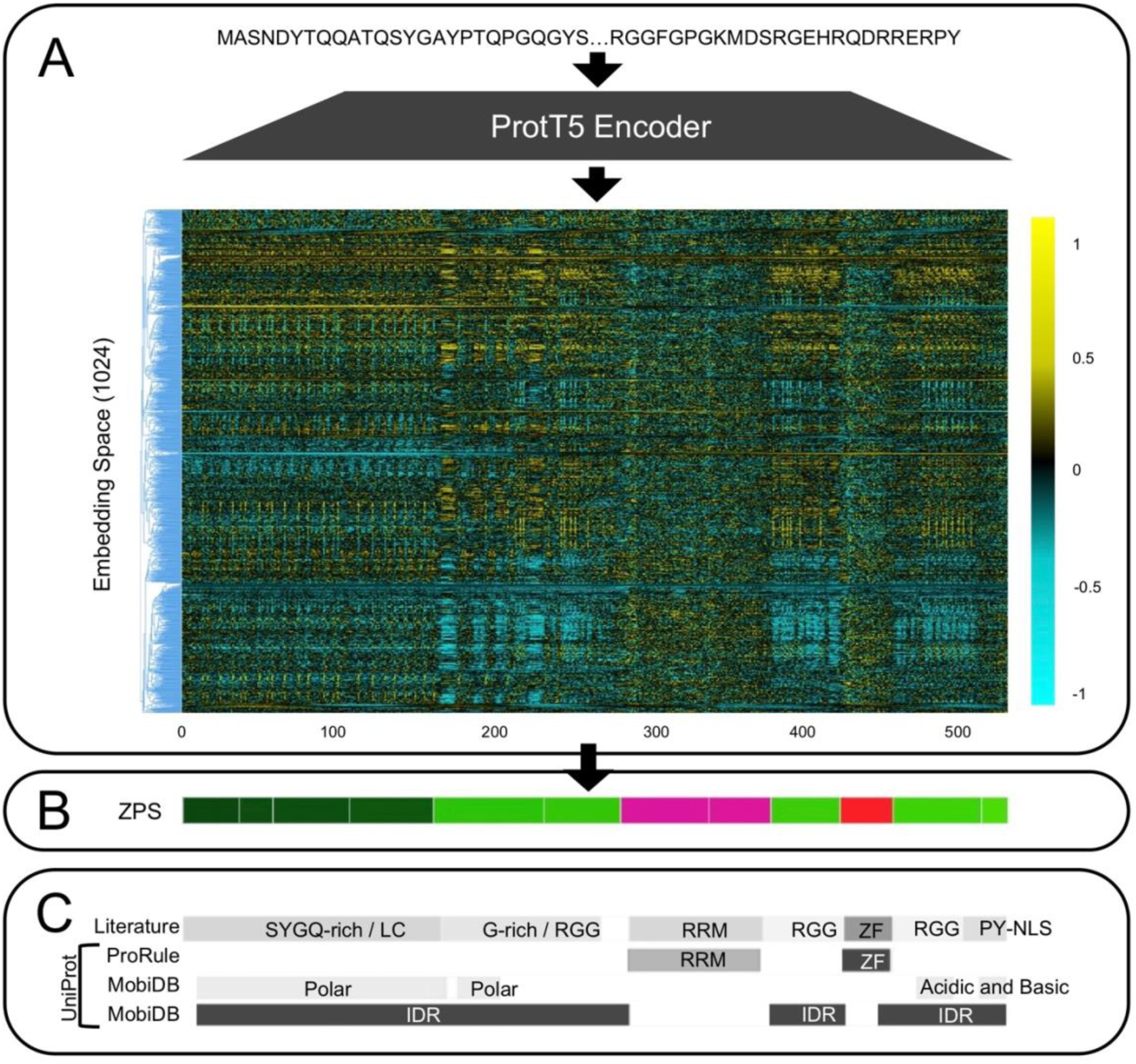
Zero-Shot Protein Segmentation Summary and Comparison to the Literature. (A) A per-residue protein embedding from ProtT5 for human protein FUS (UniProt ID: P35637). The embedding is visualized as a heatmap where the x-axis represents each amino acid along the proteins sequence (which is maintained across the panels in this Figure) and the y-axis is the ProtT5 embedding space. The embedding space is ordered by hierarchical agglomerate clustering to make patterns in the embedding space more visible. (B) Segments of FUS found using Zero-shot Protein Segmentation (ZPS). The colour of each protein segment was obtained by reducing 1024-dimensional segment embeddings to 3-dimensions, which were scaled to RGB colours (see Methods). (C) FUS annotations from the literature (see Methods) and UniProt. MobiDB and ProRule annotations were retrieved from UniProt.

We noted that in addition to known boundaries, the embedding also appears to contain higher resolution information than its annotations (Fig 1). For example, the RNA Recognition Motif (RRM) domain (folded) is broken into two segments by ZPS. Generally, RRM domains have the structure **β_1_**α_1_β_2_**β_3_**α_2_β_4_ with an RNP2 motif in **β_1_** and an RNP1 motif in **β_3_** [31] which align with the 2 faint vertical bands within the embedding of the RRM domain (amino acids 287-293 and 333-341, Fig 1A). Each of the two ZPS segments within the RRM domain begins with one of these motifs, suggesting that the pLM is distinguishing a real functional or structural boundary within the RRM domain. Similarly, within the IDRs, arginine-glycine-glycine (RGG) regions are defined by multiple short repeating motifs of arginine and glycine and are associated with specific biological functions [32]. In the RGG regions of the FUS embedding, the bold vertical lines are the arginines and the thicker vertical bands are the glycines, which clearly distinguishes these segments (amino acids 161-279, 376-419, 453-509, Fig 1A) from other IDRs. The increased magnitude in the embedding for these specific residues in the RGG regions suggests the pLM recognizes their significance in these regions of the protein. These results are consistent with previous observations that pLMs learn key functional residues and regions within proteins [15,22,33] and indicates that this observation applies to both folded domains and IDRs.

For a more compact and interpretable visualization of the high-dimensional segment embeddings, we display each segment as a block of colour along the linear protein sequence (Fig 1B), as commonly seen in the molecular biology literature [29,30,32,34]. We achieve this by averaging the per-residue embeddings over the sequence length of a ZPS protein segment to obtain a “segment embedding” (see Methods). Then the segment embedding is reduced from a 1024-dimensional space to a 3-dimensional RGB colour space to obtain a specific colour for that segment (see Methods). Just as distinct clusters in a 2-dimensional visualization of protein embeddings can be associated with biologically meaningful annotations (such as domains, evolutionary relationships, and folded structures) [15,16,19,35], distinct colours in the 3-dimensional RGB space can be associated with biologically meaningful annotations from the literature (Fig 1). For example, there are SYGQ-rich and RGG regions within the N-terminal IDR of FUS that are easily distinguishable by their segment colour and well-defined in the literature, but these are not well-characterized in UniProt (Fig 1). Despite over-segmentation relative to annotations from UniProt and the literature, segments with the same colour fall under the same annotation, suggesting that segment embeddings in the high-dimensional space can be associated with annotations (explored further in later sections).

### Segment embeddings can be used to categorize annotated regions of RNA binding proteins

To assess whether segment embeddings can be used to categorize segments across multiple proteins, we segmented well-characterized RNA binding proteins using ZPS, then compared their segments to annotations from the literature and AlphaFoldDB structures [36]. We selected the FET family (FUS, EWSR1, and TAF15), TDP-43, and HnRNPA1 which have a variety of annotated folded domains, IDRs, and regions within IDRs. The ZPS boundaries and segment embedding colours of these proteins are consistent with their folded domain and IDR sub-region annotations from the literature [29,30] (Fig 2A). Further, clustering the high-dimensional segment embeddings of these proteins separates the visually similar nuclear localization signals from RGGs (Fig 2B), consistent with the different annotations of these regions. Interestingly, the two N-terminal TDP-43 segments (Fig 2, purple) that comprise the N-terminal domain [34] cluster more closely with disordered SYGQ-rich regions than folded domains, despite AlphaFoldDB displaying a confident structure (pLDDT>90) (Fig 2C, purple).

**Fig 2.**
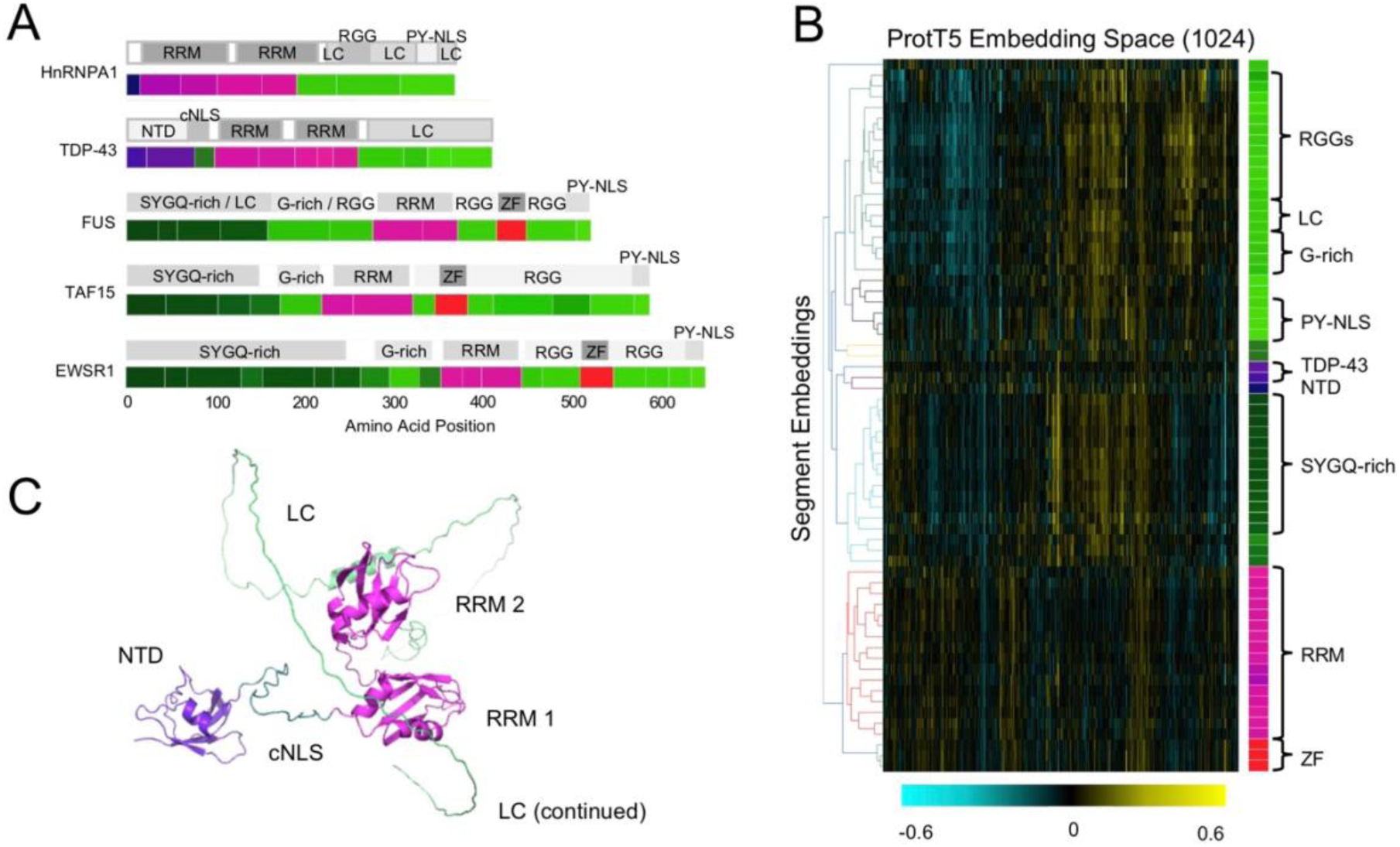
Segment Embeddings and Segment Colours of RNA-Binding Proteins. (A) Segment embedding boundaries and colours compared to annotations from the literature (see Methods) for the FET family (FUS, EWSR1 (UniProt ID: Q01844), and TAF-15 (UniProt ID: Q92804)), HnRNPA1 (UniProt ID: P09651), and TDP-43 (UniProt ID: Q13148). (B) Segment embeddings shown in panel A were ordered by hierarchical agglomerate clustering and visualized as a heatmap. Corresponding colours of the segment embeddings are shown to the right of the heatmap and labelled with annotations from the literature. (C) AlphaFoldDB structure of TDP-43 (ID: Q13148) showing segment colours. Abbreviations for folded domains labelled here include RNA Recognition Motifs (RRM), N-terminal domain (NTD), and Zinc Fingers (ZF). Abbreviations for IDR sub-regions labelled here include canonical Nuclear Localization Signal (cNLS), Low Complexity (LC) regions, PY-motif Nuclear Localization Signals (PY-NLS), and RGG (arginine-glycine-glycine motif) regions. G-rich and SYGQ-rich regions describe compositional biases that define sub-regions of IDRs.

Surprisingly, this is consistent with the literature since the stability of this structure is debated [37,38]. Taken together these results suggest that the similarity of segment embeddings can be used to categorize different folded domains and different IDR sub-regions across multiple proteins. To our knowledge, no current single bioinformatics method can simultaneously categorize both folded domains and IDRs. Additionally, the above analysis demonstrates that visualizing protein embeddings as segments in the 3-dimensional colour space offers a simple strategy for insight into unsupervised pLM embeddings.

### Zero-shot protein segmentation can reproduce the boundaries of UniProt annotations from the human proteome

To quantitatively assess the boundaries predicted by ZPS, we systematically compared segment boundaries to annotated segments in the human proteome from UniProt. We compare ZPS to Pfam [1] and PrositeScan [2], which are supervised bioinformatics tools that primarily identify folded domains in proteins. We also compare ZPS to fLPS2 [12] and Chi-Score Analysis [13], which are unsupervised bioinformatics tools that segment proteins. Additionally, we distinguish between UniProt, MobiDB, and ProRule, where MobiDB and ProRule are separate databases that are included in UniProt. MobiDB contains IDR and compositional biases annotations while ProRule contains domain annotations.

First, we evaluated this approach as a segmentation problem using Intersection over Union (IoU), a widely used measure of segmentation agreement from image analysis [25,39]. A measure similar to IoU was previously proposed for assessing protein secondary structure prediction of proteins [40]. We find that when evaluated on all human proteome annotations in UniProt (including both MobiDB and ProRule), ZPS outperforms both the supervised and unsupervised bioinformatics approaches on average IoU and recall (Table 1). This is consistent with our results for FUS and other proteins (Figs 1 and 2), where ZPS can identify the boundaries of both folded domains and IDRs. Since we have previously observed over-segmentation relative to annotations from UniProt and the literature (Figs 1 and 2), we also report metrics for over-segmentation corrected ZPS (see Methods), which improves precision and reduces recall (Table 1). We find that ZPS outperforms both supervised and unsupervised bioinformatics approaches on average IoU and recall for annotations from MobiDB (Table 1). Finally, for ProRule annotations we find that ZPS performance falls between supervised and unsupervised bioinformatics approaches, where as expected, supervised bioinformatics approaches trained on domains have the best performance (Table 1).

**Table 1.** Segmentation Evaluation for Human Proteome Segment Annotations from UniProt.

|  | All UniProt (N=91k) |  |  |  | MobiDB (N=23k) |  |  |  | ProRule (N=22k) |  |  |  |
| --- | --- | --- | --- | --- | --- | --- | --- | --- | --- | --- | --- | --- |
|  | Predicted Segments | Average IoU | Precision | Recall | Predicted Segments | Average IoU | Precision | Recall | Predicted Segments | Average IoU | Precision | Recall |
| **Supervised** |  |  |  |  |  |  |  |  |  |  |  |  |
| Pfam | 152k | 0.436 | 0.264 | 0.456 | 74k | 0.103 | 0.025 | 0.080 | 97k | 0.765 | 0.195 | 0.872 |
| Prosite Scan | 32k | 0.357 | **0.830** | 0.347 | 18k | 0.128 | 0.132 | 0.108 | 26k | **0.877** | **0.725** | **0.897** |
| **Unsupervised** |  |  |  |  |  |  |  |  |  |  |  |  |
| fLPS2 | 151k | 0.367 | 0.177 | 0.303 | 95k | 0.409 | 0.088 | 0.362 | 84k | 0.316 | 0.058 | 0.223 |
| fLPS2 (low complexity) | 57k | 0.155 | 0.199 | 0.128 | 45k | 0.344 | **0.165** | 0.327 | 33k | 0.080 | 0.038 | 0.057 |
| Chi-Score Analysis | 554k | 0.345 | 0.039 | 0.240 | 316k | 0.434 | 0.029 | 0.393 | 310k | 0.282 | 0.009 | 0.126 |
| Chi-Score (filtered) | 81k | 0.380 | 0.346 | 0.333 | 51k | 0.396 | 0.160 | 0.357 | 43k | 0.337 | 0.133 | 0.265 |
| **Our Method** |  |  |  |  |  |  |  |  |  |  |  |  |
| ZPS | 253k | **0.532** | 0.189 | **0.538** | 152k | **0.587** | 0.096 | **0.635** | 144k | 0.541 | 0.080 | 0.530 |
| ZPS (corrected) | 164k | 0.516 | 0.257 | 0.482 | 98k | 0.534 | 0.121 | 0.519 | 93k | 0.549 | 0.125 | 0.533 |

We report the number of predicted segments and Intersection over Union (IoU) metrics for established bioinformatics tools and ZPS. ZPS (corrected) uses a simple correction for over-segmentation (see Methods). The best performance for each measure is shown in **bold.** N shows the number of annotated segments in the dataset. We use an IoU threshold of 0.5 to define precision and recall.

In addition to IoU, we assessed the accuracy of boundaries by calculating the distance in amino acids between the boundaries of annotated segment and predicted segment pairs (see Methods). Overall, we found that ∼40-50% of ZPS boundaries are less than 10 amino acids from a known boundary (Table 2). This demonstrates that on average, ZPS can accurately predict about half the boundaries of known segment annotations. Furthermore, ZPS outperforms supervised and unsupervised bioinformatics tools by about 10-30% when we consider all human proteome annotations in UniProt or only this subset from MobiDB. As expected, Prosite Scan and Pfam perform best on the ProRule annotations from UniProt. Nevertheless, a change point analysis on ProtT5 embeddings can consistently outperform unsupervised bioinformatics tools developed with specific biological knowledge for IDRs and compositional biases. Furthermore, unlike other methods, our approach shows reasonable performance on both folded domains and IDRs in the human proteome.

**Table 2.**
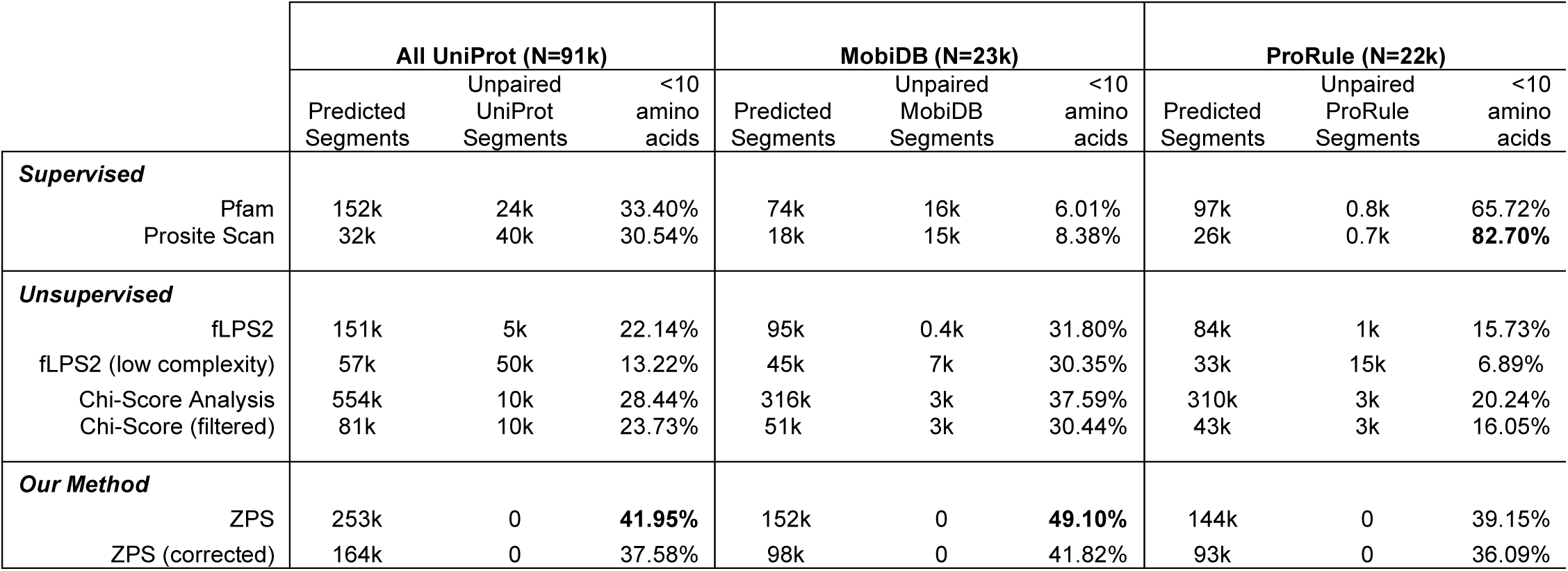
Boundary Evaluation for Human Proteome Segment Annotations from UniProt.

|  | All UniProt (N=91k) |  |  | MobiDB (N=23k) |  |  | ProRule (N=22k) |  |  |
| --- | --- | --- | --- | --- | --- | --- | --- | --- | --- |
|  | Predicted Segments | Unpaired UniProt Segments | <10 amino acids | Predicted Segments | Unpaired MobiDB Segments | <10 amino acids | Predicted Segments | Unpaired ProRule Segments | <10 amino acids |
| **Supervised** |  |  |  |  |  |  |  |  |  |
| Pfam | 152k | 24k | 33.40% | 74k | 16k | 6.01% | 97k | 0.8k | 65.72% |
| Prosit Scan | 32k | 40k | 30.54% | 18k | 15k | 8.38% | 26k | 0.7k | **82.70%** |
| **Unsupervised** |  |  |  |  |  |  |  |  |  |
| fLPS2 | 151k | 5k | 22.14% | 95k | 0.4k | 31.80% | 84k | 1k | 15.73% |
| fLPS2 (low complexity) | 57k | 50k | 13.22% | 45k | 7k | 30.35% | 33k | 15k | 6.89% |
| Chi-Score Analysis | 554k | 10k | 28.44% | 316k | 3k | 37.59% | 310k | 3k | 20.24% |
| Chi-Score (filtered) | 81k | 10k | 23.73% | 51k | 3k | 30.44% | 43k | 3k | 16.05% |
| **Our Method** |  |  |  |  |  |  |  |  |  |
| ZPS | 253k | 0 | **41.95%** | 152k | 0 | **49.10%** | 144k | 0 | 39.15% |
| ZPS (corrected) | 164k | 0 | 37.58% | 98k | 0 | 41.82% | 93k | 0 | 36.09% |

We report the number of predicted segments, the number of annotated segments without an overlapping predicted segment (Unpaired Segments), and the percentage of annotated boundaries with a predicted boundary less than 10 amino acids away. ZPS (corrected) uses a simple correction for over-segmentation (see Methods). The best performance for each measure is shown in **bold.** N shows the number of annotated segments in the dataset.

### Segment embeddings can distinguish folded domains from intrinsically disordered regions and 5 different compositional biases

Our results for well-characterized RNA binding proteins (Fig 2) suggested that segment embeddings can distinguish different types of regions within proteins. So, we tested whether segment embeddings can be used to distinguish protein segments with different biological properties associated with function, such as compositional biases and whether the protein segment can fold independently. Protein segments defined by ZPS were labelled by transferring annotations from UniProt with the highest IoU (see Methods) and then compared to their 1-nearest neighbour (1-nn) in the high dimensional segment embedding space. Here, we find that MobiDB IDRs and ProRule domains are clearly separated in the segment embedding space. The 1-nn accuracy (see Methods) for ProRule domains and MobiDB IDRs are 0.985 +/-0.002 and 0.979 +/- 0.002, respectively (Fig 3A). This is as expected since IDRs and domains can be distinguished based on amino acid composition, which we confirmed using one hot encoding embeddings of these segments (0.895 +/-0.004 and 0.904 +/-0.004 for ProRule domains and MobiDB IDRs, respectively). Similarly, segment embeddings can distinguish the 5 different types of compositional bias annotations, where the 1-nn accuracies range from 0.687 to 0.820 with confidence intervals ranging from +/-0.012 to +/-0.059 (Fig 3B). By reducing the dimensionality of the segment embeddings using a UMAP, we can visualize segment embeddings in a scatter plot. This shows that folded domains and IDRs are clearly separable (Fig 3C). The compositional biases visualization shows more overlap between protein segments with different compositional biases, but still reveals patterns associated with their compositional bias (Fig 3D). This demonstrates that segment embeddings contain sufficient information to categorize structural state and compositional biases.

**Fig 3.**
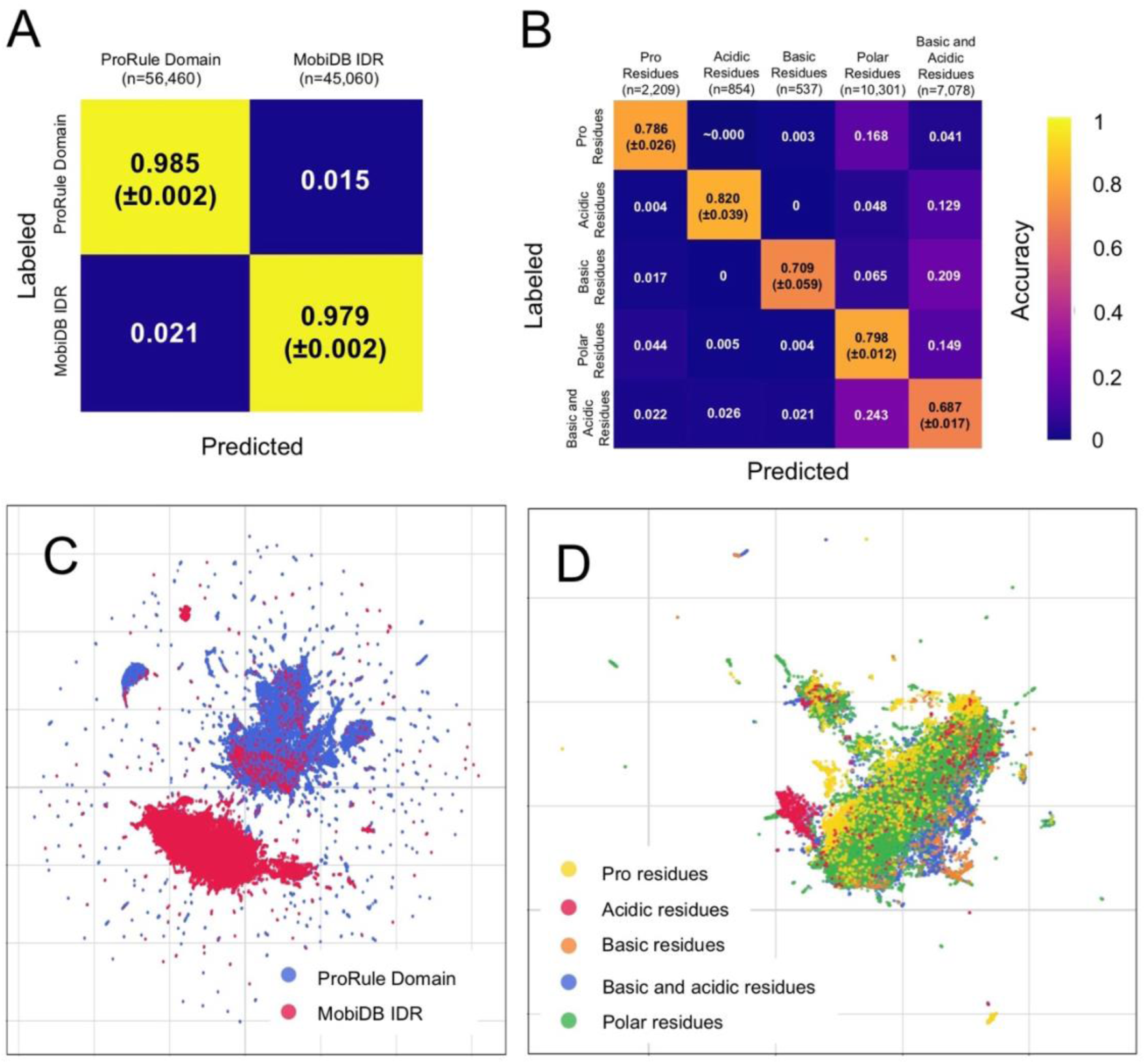
Segment Embedding Evaluation and Visualization of IDRs, Domains, and Compositional Biases. (A, B) Normalized confusion matrices for 1-nn assessment for (A) ProRule Domain compared to MobiDB Disorder Consensus and (B) MobiDB compositional biases. In the confusion matrix, we report 1-nn accuracy with the binomial proportion confidence interval shown below in brackets for true positive predictions (n is the number of ZPS segments). (C, D) Shows 2-dimensional UMAPs of segment embeddings labelled with (C) ProRule domains compared to MobiDB disorder consensus and (D) MobiDB compositional biases. Each point in the UMAP is a protein segment, and each segment shown in the UMAP was used in the respective 1-nn assessment.

### Segment embeddings can distinguish different types of protein domains and sub-domains

We next tested whether segment embeddings can distinguish different types of protein domains. Here, we transfer ProRule annotations from UniProt to segment embeddings and analyze the top 20 most common domain annotations amongst the segment embeddings using 1-nn. We find different types of domains are easily distinguishable and have 1-nn accuracies that range from 0.935 to 0.997 with a confidence interval ranging from +/-0.001 to +/-0.036 (Fig 4A). To confirm that segment embeddings from the T5 model are more informative than embeddings of simple amino acid sequences, we compare our results to 3-mer embeddings (see Methods) of the same protein segments. Segment embeddings show an average of 0.241 increase compared to 3-mer embeddings, where their 1-nn accuracies range from 0.435 to 0.992 (+/-0.009 to +/- 0.078) (S1 Table).

**Fig 4.**
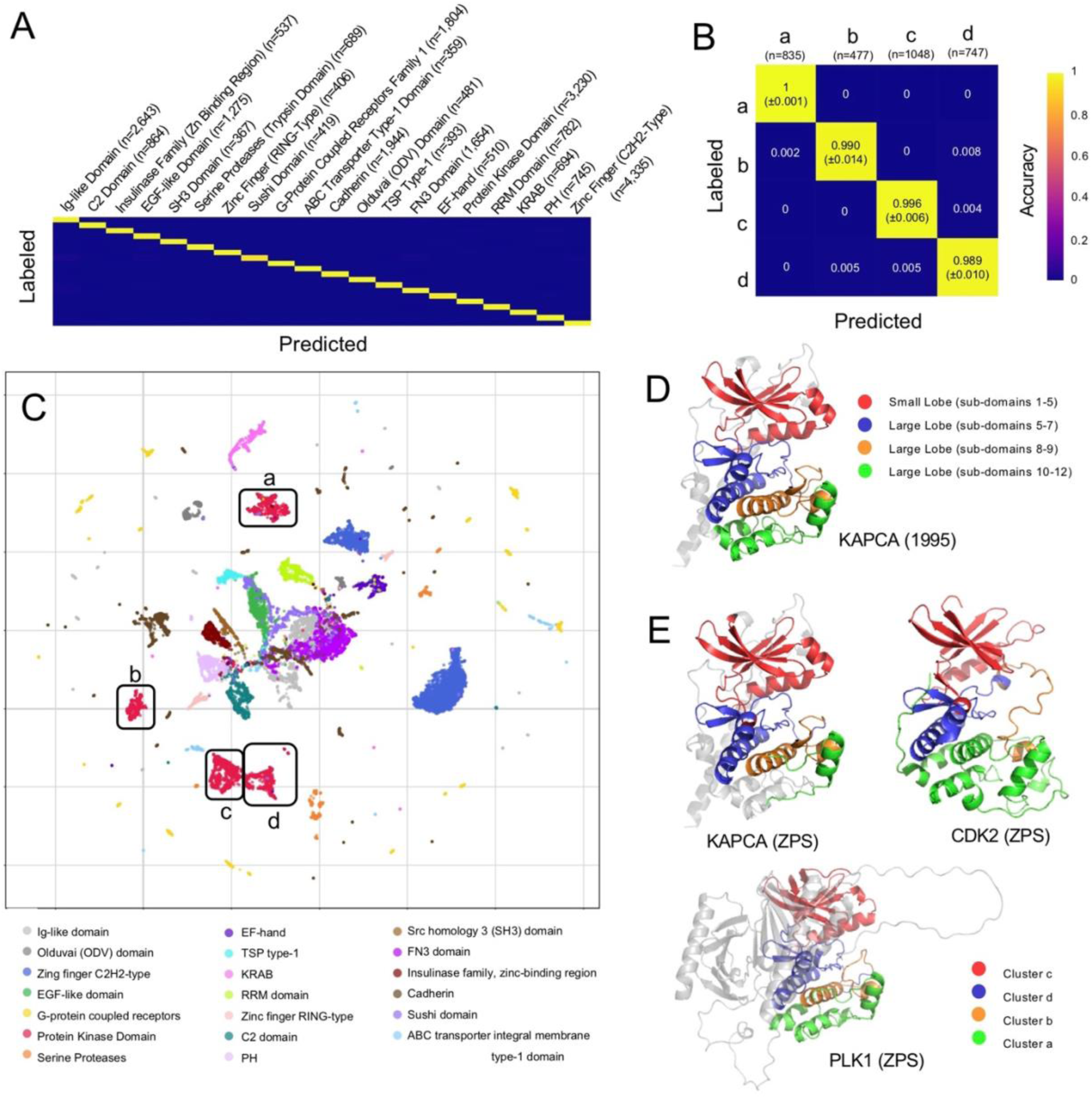
Segment Embedding Evaluation and Visualization Domain Types and Sub-Domains. (A, B) Normalized confusion matrices for 1-nn assessment (n is the number of ZPS segments). (A) Top 20 most commonly occurring ProRule domains in the ZPS segments and (B) shows clusters a, b, c, and d of the protein kinase domain (as labelled in panel C). In (B) we report 1-nn accuracy with the binomial proportion confidence interval shown below in brackets along the diagonal (S1 Table). (C) A 2-dimensional UMAP of segment embeddings coloured by the type of domain the segment overlaps with. This shows clusters a, b, c, and d of the protein kinase domain segments. Each of the protein segments shown here were used in the 1-nn assessment. (D) The AlphaFoldDB structure of KAPCA (UniProt ID: P17612) with the protein kinase domain shown in colour, with sub-domains 1-5 (red), 5-7 (blue), 8-9 (orange), and 10-11(green). Sub-domains 1-5 are the small lobe and 5-11 are the large lobe, as defined in [41]. (E) AlphaFoldDB structures of KAPCA, CDK2 (UniProt ID: P24941), and PLK1 (UniProt ID: P53350), where colours are defined by clusters of protein kinase segments, including cluster c (red), cluster d (blue), cluster b (orange), and cluster a (green).

We visualized segment embeddings with different domain annotations using UMAP, where we noted some domains organized into multiple clusters, such as the protein kinase domain (Fig 4C and S2 Table). First, we established that these clusters exist in the high dimensional space by performing the same 1-nn analysis and found the 1-nn accuracies range from 0.989 to 1 (Fig 4B). Once again, the T5 segment embeddings are more informative than 3-mer embeddings with 1-nn accuracies ranging from 0.849 to 0.948 (+/-0.027 to +/-0.037). We then found the segments in these clusters can be ordered linearly within the protein kinase domain amino acid sequence, starting from the N-term with cluster c, then d, then b, and cluster a, for 425/481 (88%) protein kinase domains in the human proteome. From this, we determined that segments in clusters a, b, c, and d are associated with sub-domains of the protein kinase domain. The protein kinase domain is comprised of 12 sub-domains which can be grouped into 2 lobes, a small lobe containing sub-domains 1-5 (Fig 4D, red) and a large lobe containing domains 5-11 (Fig 4D, blue, orange and green), where sub-domain 5 spans both lobes [41]. We found that the small lobe (Fig 4D, red) corresponds to segments from cluster c (Fig 4E, red). In the large lobe, sub-domains 5, 6A, 6B, and 7, (Fig 4D, blue) identified by their structural components (a-helix D, a-helix E, and B-strands 6-9 [41]), correspond to cluster d (Fig 4E, blue). We observe slight variations with the sub-domains included in clusters a and b, most notably whether cluster a includes a few extra alpha helices (Fig 4E, CDK2) or is missing a few alpha helices (Fig 4E, PLK1). Despite these small variations in how a few short alpha helices are distributed between clusters a and b, our approach can zero-shot segment protein domains and sub-domains and distinguish them more effectively than 3-mer embeddings.

### Segment embeddings can distinguish intrinsically disordered regions

We next assessed whether segment embeddings can be used to distinguish different IDRs. This is a difficult problem for which few supervised approaches have been developed [7,11]. We selected segments that overlap with IDR annotations from MobiDB then transferred additional overlapping annotations, such as compositional biases, to define different IDRs, and then selected the top 20 most common IDR annotations (see Methods). We use this set of IDR annotations because there is a lack of consensus on IDR types and identifying a broad range of meaningful large-scale annotations remains challenging. Then we calculated the 1-nn accuracy and visualized the segment embeddings of the IDRs using UMAP (Fig 5). We observe a wide range of 1-nn accuracies with an average of 0.650 (S1 Table), which we believe is due, at least in part, to the broader definition of IDR annotations compared to folded domain annotations. For example, we have labels with non-exclusive meanings such as Collagen-Related and Triple Helical Region as well as COILED and HELIX [42,43].

**Fig 5.**
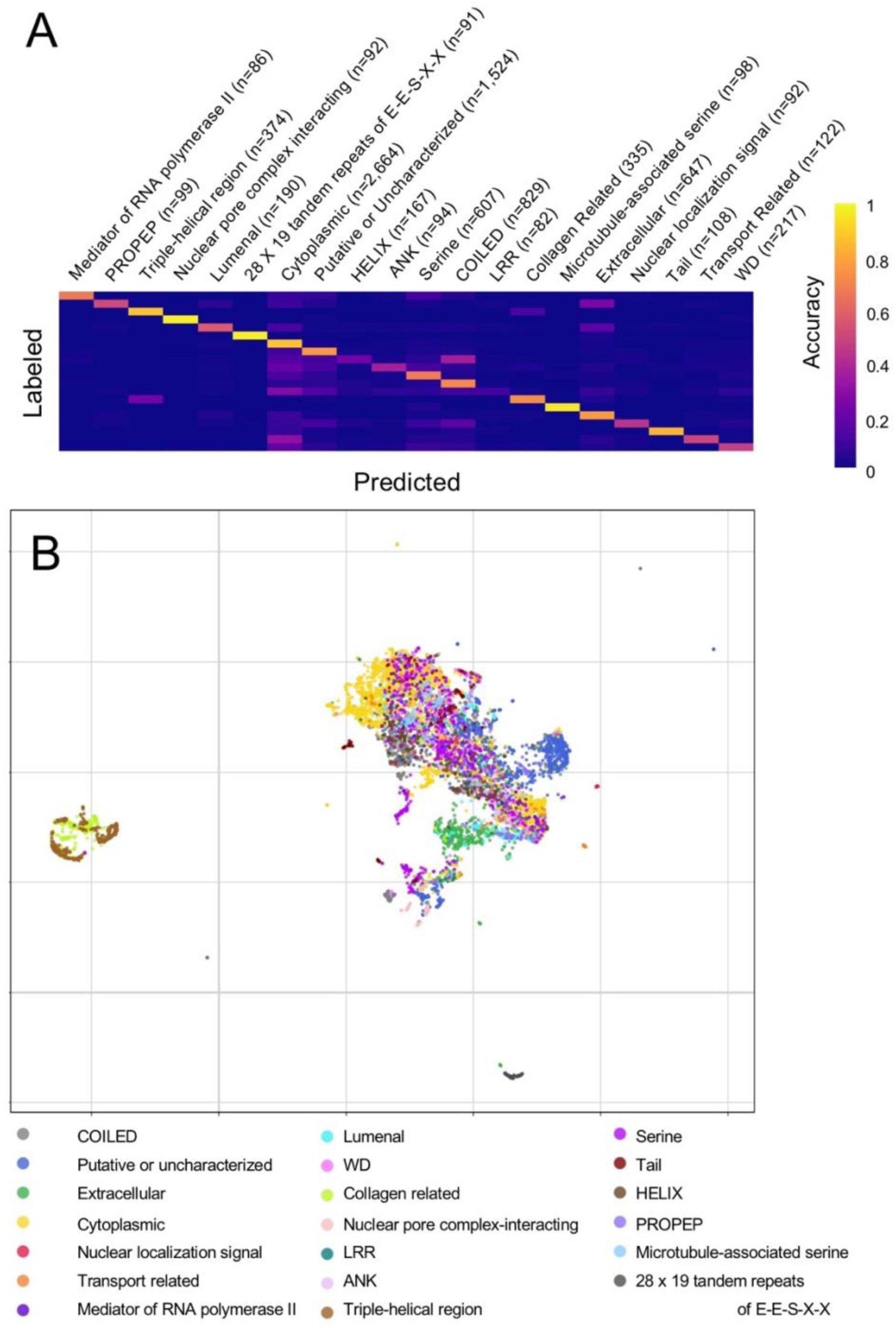
Segment Embedding Evaluation and Visualization of IDRs. (A) The normalized confusion matrix for 1-nn assessment of the top 20 most common annotations that overlap with IDRs defined by MobiDB. n is the number of ZPS segments with each annotation. (B) UMAP of the segments used in the 1-nn assessment labelled with the IDR types.

Nevertheless, we see a clear improvement over random (average improvement of 0.561) and 3-mer embeddings of the same segments (average improvement 0.339) (S1 Table). This demonstrates that segment embeddings can be used to distinguish different IDRs.

### Discovery of unannotated mitochondrial targeting signals in the human proteome

To investigate whether the segment embeddings can be used to make predictions about unannotated protein segments, we used Leiden clustering [44,45] to identify clusters of unannotated segments in the human proteome (see Methods). We then projected all human protein segment embeddings to a 2-dimensions space using UMAP to visualize clusters of unannotated segments in the context of the whole human proteome. After correction for over-segmentation (See Methods) we find 51 unannotated segments that are separate from the majority of the human proteome (Fig 6A, S3 Table). We found that there are 609 annotated segments that appear near this cluster on the UMAP (S3 Table), which overlap with 435/495 (87.8%) of the annotated mitochondrion targeting signals in the human proteome from UniProt. Consistent with this, since typical mitochondrial targeting signals are located at the N-terminus of the protein sequence [46], 644/660 (97.6%) of these segments (annotated and unannotated) begin at the N-terminus of the protein. Further, 625/648 (96.5%, 7 unmapped proteins) of the proteins containing these segments localize to the mitochondria (p=0), including 44/48 (91.7%, 3 unmapped proteins) proteins with unannotated segments (p=8.98E-39). Interestingly, UniProt includes 58 mitochondrion targeting signals of unknown length in the human proteome that begin at the N-terminus of the protein, of which 39 (76.5%) are found in this cluster of 51 unannotated segments. This evidence strongly suggests that the remaining 12 unannotated segments in this cluster (Fig 6A, cyan) are undiscovered mitochondrion targeting signals, and we can use this information to determine the size of the 39 mitochondrion targeting signals of unknown length.

**Fig 6.**
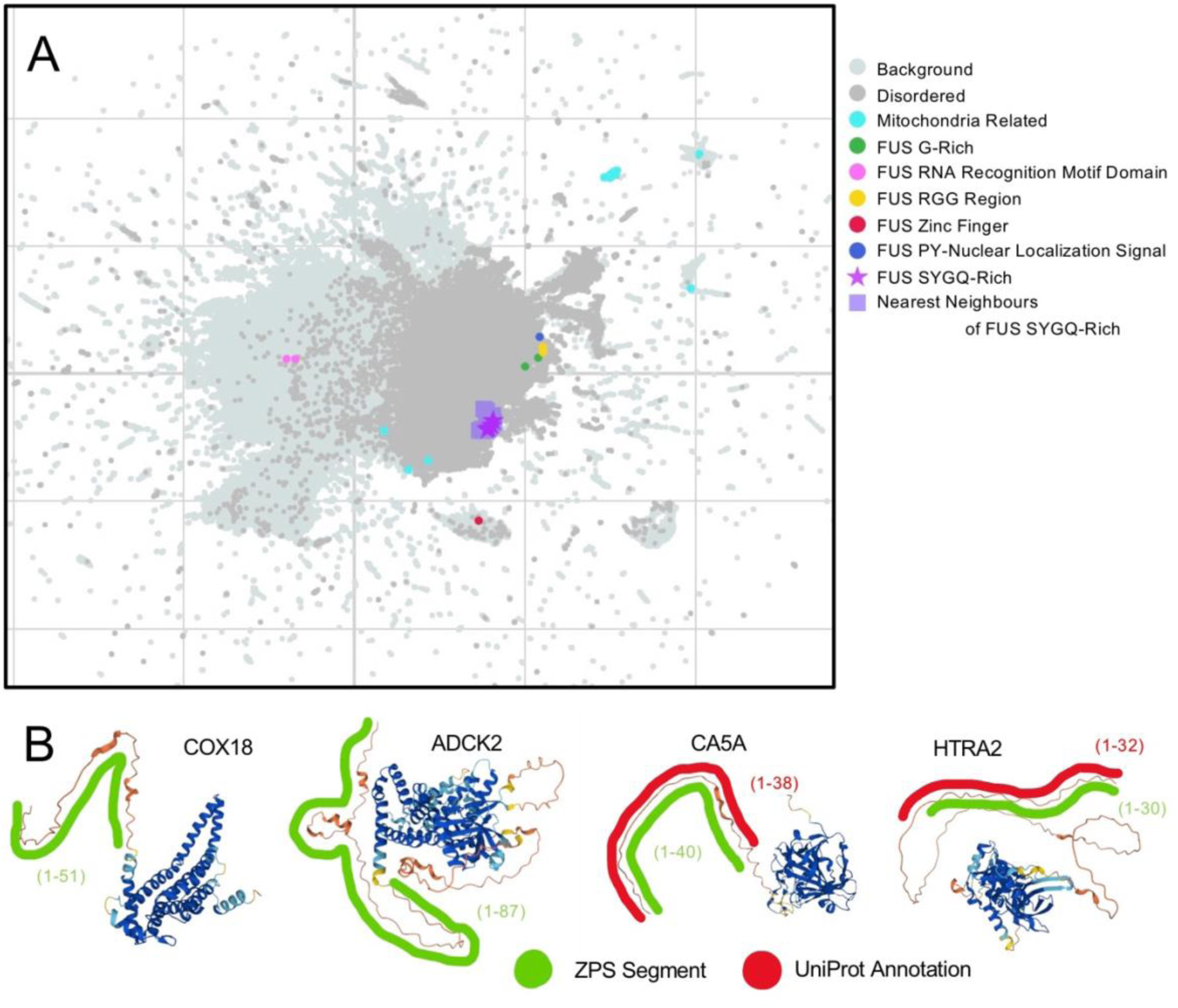
Identification of Mitochondrion-Related Cluster and SYQG-Rich Prion-Like Domains Similar to FUS. (A) UMAP of segment embeddings of the entire human proteome. Disordered segments (defined by MobiDB) are shown in a warmer shade of grey to give context to the UMAP. Mitochondria Related cluster (cyan) was defined by Leiden clustering of unannotated protein segments (see Methods). Segment embeddings of FUS are shown along with the top 10 nearest-neighbours to the segment embeddings of FUS’s SYGQ-rich regions. (B) AlphaFoldDB structures of COX18 (UniProt ID: Q8N8Q8), ADCK2 (UniProt ID: Q7Z695), CA5A (UniProt ID: P35218), and HTRA2 (UniProt ID: O43464) colored by pLDDT confidence score (very low in orange, low in yellow, high in cyan, and very high in blue), showing UniProt mitochondrion targeting signal annotations (red) and ZPS segments (green). COX18 has a UniProt mitochondrion targeting signal annotation of unknown length starting at the N-terminus and ADCK2 has no UniProt mitochondrion targeting signal annotation but is known to localize to the mitochondrion [47].

To test whether segment embeddings associated with this unannotated segment cluster (Fig 6A, cyan) are simply the N-terminal IDRs of mitochondrial proteins instead of actual mitochondrion targeting signals, we compared examples of these segments to predicted structures from AlphaFoldDB and annoated mitochondrial targeting signals from UniProt. In some cases, such as CA5A, the boundaries of UniProt annotations of mitochondrial targeting signals (Fig 6B, red) align closely with ZPS boundaries (Fig 6B green) and a region having AlphaFold pLDDT confidence scores <50 (“very low”, highly correlated with structural disorder, Fig 6B, orange). However, in other cases such as HTRA2, the UniProt mitochondrion targeting signal occupies only the first 30 residues (Fig 6B, red) with the ZPS boundaries matching the UniProt annotation (Fig 6B, green), while the very low confidence region spans the first 131 residues (Fig 6B, orange). This use of ZPS and segment embeddings to annotate mitochondrion localization signals provides evidence that pLM embeddings can provide valuable insight without training or fine-tuning.

### Identifying similar IDRs to FUS’s SYGQ-rich region using the k-nearest neighbours of segment embeddings

We chose to investigate the 4 SYGQ-rich segments in FUS because these segments represent “prion-like domain” elements of IDRs with well-studied role in forming condensates, such as in response DNA damage stress, or in forming aggregates, such as in amyotrophic lateral sclerosis and frontotemporal dementia [48–50]. In general, prion-like domains have a strong compositional bias and can be identified from amino acid sequences using Prion-Like Amino Acid Composition (PLAAC) scores [51].

We identified the 10 nearest-neighbours for each of the 4 segments of FUS’s SYGQ-rich prion-like domain in the human proteome. After removing duplicates and joining adjacent segments (see Methods), we have 14 segments from 11 proteins (S4 Table). On average the amino acid content of these segments is 56% SYGQ, and 771/943 (81.8%) amino acids are predicted to be in a prion-like domain by PLAAC (Fig 7 shows 4 examples). Encouragingly, the SYGQ-rich regions of all 3 FET proteins (FUS, EWSR1, and TAF-15) were included in these segments, given the known conservation of these prion-like domains across the FET family [30,52,53]. Despite visually similar sequence composition in these amino acid sequences (Fig 7), performing a default protein BLAST [54] in the human proteome using FUS’s SYGQ-rich prion-like domain only retrieves anomalous fusion proteins and isoforms of FUS and EWSR1 – the search does not identify any other proteins.

**Fig 7.**
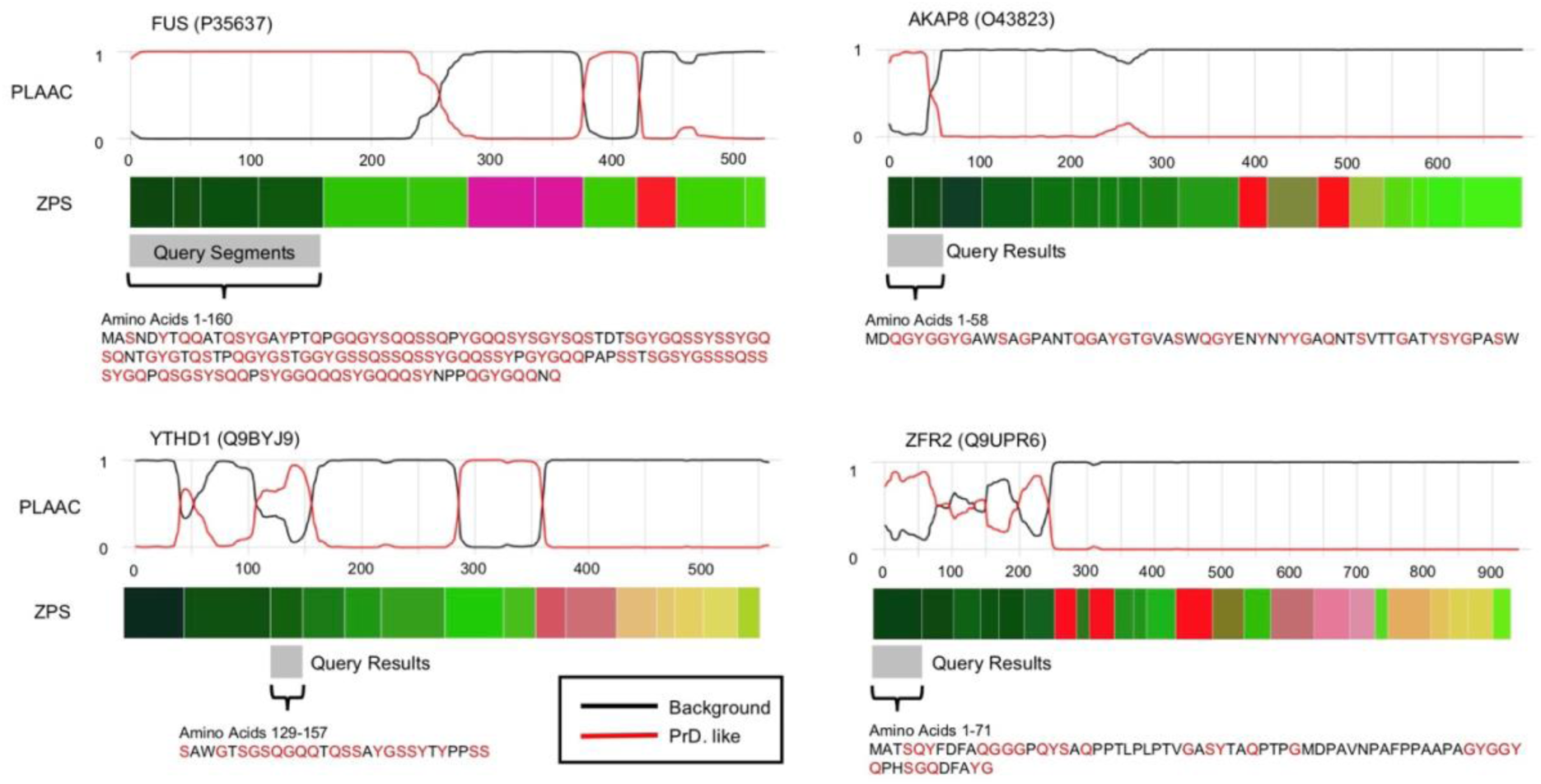
Zero-Shot Protein Segmentation of Proteins with Similar Segments to FUS’s SYGQ-Rich Region and PLAAC scores. Prion-Like Amino Acid Content (PLAAC) scores adapted from the web tool described in [51], segment embedding visualization (ZPS) as shown in Fig 1 and Fig 2, and the amino acid sequence of the query segments from FUS and query result segments (S, Y, G, and Q shown in red).

Interestingly, in addition to the FET family, we identified 8 other proteins with similar segment embeddings to FUS’s SYGQ-rich regions. Encouragingly, the segments in 3 of the 8 proteins, YTHD1, YTHD3, and AKAP8, have been reported to form condensates and bind RNA, just as the FET family’s SYGQ-rich regions can. Similar to FUS, the N-terminal IDR in YTHD1 and YTHD3 is required to form condensates with RNA in the cytosol in response to stress [55,56]. Also, similar to FUS, AKAP8 forms condensates in the nucleus and binds RNA to regulate splicing, and condensate formation is required for RNA regulation [57]. While other elements of AKAP8 have been identified to be required for condensation [57], we predict that the prion-like domain segment (Fig 7) makes a significant contribution to condensation. We observed other prion-like domains predicted by PLAAC that are not nearest-neighbours of FUS (Fig 7). However, in FUS these segments have a different compositional bias, G-rich or RGG (which also has high G content), that is also known to be enriched in prion-like domains [51]. This suggests we can distinguish prion-like domains by their compositional biases, beyond the binary classification provided by PLAAC. Taken together, our analysis suggests that the most similar segment embeddings to those of FUS’s SYGQ-rich prion-like domain are other prion-like domains with similar compositional biases that cannot be found using BLAST, or uniquely identified by PLAAC.

## Discussion

Here we show that embeddings from ProtT5 can be used to predict the boundaries of protein regions and categorize protein segments without training or fine-tuning any parameters. We predicted the boundaries of annotations from UniProt more effectively than established bioinformatics tools and demonstrated that segment embeddings are more informative than k-mer embeddings of the same sub-sequences at categorizing protein domains, sub-domains, and types of IDRs. We then used these approaches to identify unannotated mitochondria localization signals and SYGQ-rich prion-like domains in the human proteome. Additionally, we present a novel approach for visualizing pLM embeddings using protein segments and colours.

The boundaries defined by ZPS over-segment proteins relative to annotations on UniProt and the literature (Fig 1, 2, 4). While this leads to poorer performance when benchmarking against annotations from UniProt, when we investigated well-annotated proteins, we found that the over-segmentation captured meaningful biology such as motifs in structured domains and IDRs (Fig 1, RRM and RGG) as well as sub-domains (Fig 4). We also observe over-segmentation relative to IDR annotations where some segments align with different annotations (Fig 1, C-term RGG and PY-NLS), while other instances of over-segmentation (Fig 1, N-term multiple SYGQ-rich segments) remain unexplained. Given the motifs and sub-domains identified by the increased segmentation relative to UniProt and some literature annotations, it’s likely that this “over-segmentation” also represents a more detailed and informative annotation.

To visualize the high-dimensional segment embeddings, we represented the segments as colours along the protein sequence length. Colouring proteins by distinct annotations is widely done in the molecular biology literature to help understand complex proteins [29,30,32,34]. We found that for well-characterized RNA-binding proteins, we could reproduce these diagrams using boundaries defined by ZPS and colours resulting from a dimensionality reduction on segment embeddings. We believe that these visualizations will be a useful tool for biologists to visualize and compare the functional organization of proteins that they study. Further, we believe that converting embeddings from unsupervised language models to RGB-colour space represents a new way to visualize the information learned during pre-training that can be applied to any language model (A. G. Sangster, Micaela Consens, A. M. Moses, in preparation).

Our approach demonstrates an advantage over classical bioinformatics approaches based on sequence similarity, such as HMMs and alignments [2,54,58], because it does not rely on positional evolutionary conservation of primary amino acid sequence.

Consistent with this, ZPS can identify the boundaries of IDRs and regions with compositional biases (Table 1 and 2), which have poor positional conservation at the primary amino acid level but have meaningful conservation of amino acid properties distributed across their sequence [8,9]. Furthermore, these types of protein segments can be categorized using segment embeddings (Fig 2, 3, 5). Additionally, we can use segment embeddings to identify IDR protein segments that are believed to have similar functions, such as mitochondrial targeting signals and SYGQ-rich prion-like domains that drive condensation.

Finally, unlike other deep learning methods that use supervised learning, fine-tuning, or transfer learning, our approach is zero-shot. Generalizability is therefore less impacted by the content or availability of training data. This enables us to identify protein regions for which there are too few known examples to train a supervised model, such as SYGQ-rich prion-like domains. In principle, this means we can discover novel categorizations of protein regions that do not have any current characterized examples. However, task-specific fine-tuning on per-residue embeddings would likely lead to improved performance for identifying boundaries and categorizations of known regions for which there is sufficient training data. This has been demonstrated recently for genome language models that were fine-tuned for genome segmentation [59].

Looking forward, we anticipate that the continuing developments in image segmentation and deep learning can be applied to make further advancements in protein annotation and genomics. For example, leveraging the drop loss function presented in zero-shot image segmentation [25], which allows for novel object discovery, can enable pLMs to discover overlapping, novel, and biologically meaningful regions in proteins. Our work is a first step towards integrating the advancements of zero-shot image segmentation and protein language models for understanding functional organization within proteins.

## Methods

### RNA-binding protein annotations from the literature

Annotations from the literature are shown in Figs 1 and 2 for RNA binding proteins, including FUS (UniProt ID: P35637), EWSR1 (UniProt ID: Q01844), TAF-15 (UniProt ID: Q92804), hnRNPA1 (UniProt ID: P09651), and TDP-43 (UniProt ID: Q13148). We show annotations along the protein’s sequence as described in the literature [29,30,34].

### UniProt protein sequences and annotation data

These experiments use protein sequences and annotations for the human proteome from UniProtKB/Swiss-Prot (UniProt) [27] (available on Zenodo). The sequences on UniProt have reviewed annotations from a variety of sources including bioinformatics tools, academic literature, and other databases. Here “annotations” only refer to annotations that are associated with specific amino acids in the protein (available on Zenodo). We removed annotations that include non-numeric characters in their sequence positions, such as “1-?”, “100-?150” and “<1-100”, as it indicated uncertainty in the position or length of the sequence in the annotation. For evaluating protein segmentation, we use annotations from UniProt with any evidence level, including those from sequence analysis, prediction, curation, citation, MobiDB [28], and ProRule, a database provided by Prosite [2]. To assess performance on specific types of protein segments, we include a partition of UniProt that only includes IDR consensus predictions and compositional bias annotations from MobiDB. Additionally, we use a partition of UniProt that only includes domains annotated by ProRule.

### Zero-shot protein segmentation (ZPS)

We use the ProtT5 [14] encoder to generate protein embeddings for the whole human proteome for sequences less than 8k amino acids in length. Here we use “embedding” to refer to the output of the ProtT5 encoder. The ProtT5 embedding is a 2-dimensional matrix where one dimension is the length of the protein and the other is the embedding space of length 1024 (L×1024). Next, a change point analysis is performed on each ProtT5 embedding (see below for Change Point Analysis). This defines the boundaries between protein segments. For example, if the boundaries are a, b, and c in a protein of length 100, the segments are (0, a), (a, b), (b, c), and (c, 100). These ranges include the first position and go up to but not including the second, as conventionally seen in zero-based indexing. These segments are defined without fine-tuning or training any parameters.

### Change point analysis

A change point analysis is commonly used to detect changes in a signal or time series [60]. We selected this method to perform zero-shot protein segmentation because it does not require training parameters. We use a sliding window with a window size of 30 for our search algorithm and RBF kernel as our cost function. We chose a sliding window algorithm because this has previously been used in protein biology to identify modular organization within proteins [13]. This compares a segment of 15 amino acids to the next 15 amino acids for all amino acids in the protein. If there is a significant change between the first 15 amino acids and the next 15 amino acids, the space between them becomes a boundary that defines protein segments. See [60] for sliding window algorithm and RBF kernel function definition.

The number of boundaries predicted by the change point analysis is determined by one of two pre-selected hyperparameters: either set a threshold on the cost function that acts across the whole dataset or set a threshold on the number of boundaries for each protein. Interestingly, we found a way to avoid doing either, making our approach free from hyper-parameter selection in addition to not training or fine-tuning any parameters. Initially we set a threshold on the number of boundaries per protein based on the protein’s length, but we found that allowing more than 3 boundaries per 100 amino acids did not increase the total number of predicted boundaries. Therefore allowing 3 or more boundaries per 100 amino acids allows the change point analysis to predict the maximum number of boundaries it can find. To allow the method to function without restraint, we gave the threshold of 3 boundaries per 100 amino acids, which yields approximately 253k segments. Compared to methods we benchmark against, such as Pfam (152k segments), fLPS2 (151k segments), and Chi-Score Analysis (554k segments), 253k segments is reasonable considering Pfam and fLPS2 are tailored towards a single type of protein segment whereas we are attempting to predict any type of protein segment, so we moved forward with this set of predictions.

### Segment embeddings

We define segment embeddings by cutting the ProtT5 embedding at the boundaries defined by the change point analysis and then average pooling the segmented embedding to 1×1024. For example, if a segment spans amino acids (100, 125) the ProtT5 protein embedding would be cut at position 100 and 125 producing an embedding of size 25×1024 which would be average pooled down to 1×1024. Segment embeddings were generated without training or fine-tuning any parameters.

### K-mer embeddings

We use k-mer embeddings as a baseline comparison for the content of the segment embeddings. K-mer embeddings are defined using the same protein segments as segment embeddings. 1-mer embeddings are equivalent to one hot encoding where each dimension in the embedding represents the number of occurrences of each amino acid. 3-mer embeddings contain the number of occurrences of each permutation of 3 amino acids (AAA, AAC, AAD, …). We use overlapping k-mers and normalize k-mer embeddings before evaluations.

### Correcting for over-segmentation

After analyzing FUS and other RNA-binding proteins, it was apparent that ZPS over-segments relative to annotations on UniProt despite the meaningful biology, such as motifs, that is captured by this over-segmentation (Fig 1 and 2). We attempt to correct for over-segmentation by merging adjacent segments if they have similar segment embeddings. We measure the cosine similarity between all segments in a protein; if the segments with the highest cosine similarity are adjacent to each other in the protein sequence, then they are merged. We only correct for over-segmentation if there are 6 or more segments in a protein since this algorithm leads to poor results for proteins with a small number of segments. For example, if there are only 3 segments in the protein, then the middle segment will always get merged since it is adjacent to all the other segments of the protein, regardless of their similarity. Increasing the number of segments to 6 reduces the chance of “accidental” merging while still reducing the total number of segments by ∼100k (40%).

We also correct for over-segmentation when analyzing clusters (Fig 4 and 6). Here, we simply join adjacent protein segments that are in the same cluster.

### UMAPs

We used Uniform Manifold Approximation and Projection (UMAP) [61] to reduce the dimensionality of segment embeddings in two different instances. One is to visualize segment embeddings in a 2-dimensional scatter plot and the other is to visualize segment embeddings as 3-dimensional colours (see next section).

### Segment embeddings to RGB colours

For our visualization approach, we reduced the segment embeddings (1×1024) of the whole human proteome to 3 dimensions (1×3) using a UMAP [61]. Each of the 3 dimensions is mean-centered across the whole proteome. Then we apply a sigmoid function to each of the RGB values for all segments and multiply by 255. We do this to scale the values and change their distribution from an approximately normal distribution to a more uniform distribution. This shifts the colours from mid-tone greys to increase their vibrancy and make the colours more distinct from each other. After this step, each segment has 3 values ranging from 0-225, one for red, green, and blue, which can be directly converted to a specific RGB colour.

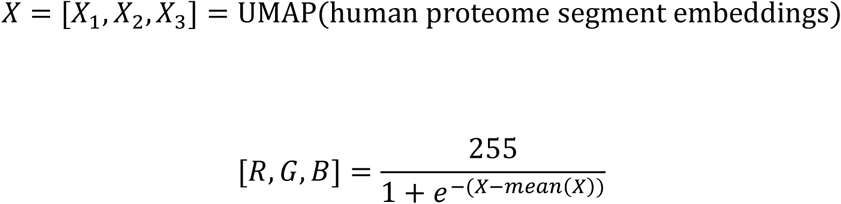

This function is applied element-wise, where X_1_ maps to R, X_2_ maps to G, and where X_3_ maps to B.

### Supervised bioinformatics approaches for protein segmentation

To our knowledge, there are not any supervised bioinformatics approaches designed for segmenting proteins that do not focus only on a specific type of protein region, such as domains. So, we use two supervised bioinformatics approaches that are designed to identify and categorize protein domains, Pfam and Prosite. We use HMMER 3.1b2 [58] to identify protein domains defined by Pfam (Pfam-A.hmm) [1] with default parameters. Additionally, we define another set of domains with Prosite Scan from ps_scan_linux_x86_elf.tar.gz (updated 2018) also using default parameters [2]. Prosite Scan is not to be confused with ProRule. While both are provided by PROSITE, Prosite Scan is an automatic domain annotation tool and ProRule is more of a “master set” of domain annotations including domains defined by manual curation. ProRule is included in UniProt.

### Unsupervised bioinformatics approaches for protein segmentation

We use unsupervised bioinformatics approaches designed to segment proteins according to changes in amino acid compositional biases as a contrast to domain annotation tools. We ran fLPS2 [12] on the whole human proteome with default parameters and parameters specifically for short low complexity compositional biases (./fLPS2 -t1e-5 -m5 -M25, see fLPS2 README for further details). fLPS2 predicts protein segments with compositional biases and can make multiple overlapping predictions; we included overlapping predictions in the evaluation. We also ran a Chi-Score Analysis [13], which predicts boundaries where there are significant changes in amino acid composition and has a built-in boundary filtering approach. We evaluated this approach with and without filtering boundaries. For Chi-Score Analysis we only included sequences less than 2k in length due to time complexity issues.

### Intersection Over Union (IoU)

The Intersection over Union (IoU) is the number of amino acids in the intersection of a pair of predicted and annotated segments divided by the number of amino acids in the union of those two segments. This produces a measurement that is relative to the size of both segments. Predicted segments include those defined by ZPS, Pfam, Prosite Scan, fLPS2, and the Chi-Score Analysis. Annotated segments include human protein annotations from UniProt that are at least 30 amino acids in length and do not span the entire length of the protein from proteins that are at least 60 amino acids in length.

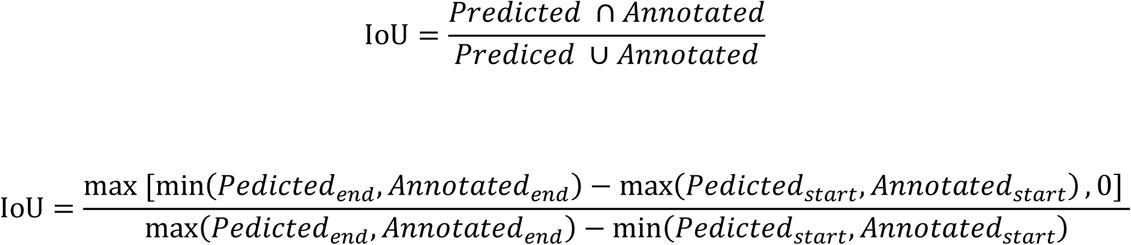

Next, we measure the average IoU. For each annotated segment we select the predicted segment with the largest IoU value as its pair, then average over all annotations. We use a threshold of 0.5 on IoU to define precision and recall. If a pair of segments has an IoU above the threshold, it is a true positive. If a predicted segment does not have a pair above the threshold, it is a false positive. If an annotated segment does not have a pair above the threshold, it is a false negative. Precision measures the proportion of predicted segments that have a paired segment above the threshold and recall measures the proportion of annotated segments that have a paired segment above the threshold.

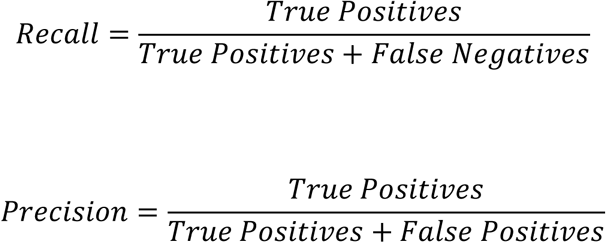

### Boundary evaluation

To evaluate the accuracy of the predicted boundaries, we select predicted and annotated segment pairs by those with the highest IoU (as described above). Then we measure the difference in amino acids at the beginning and end of the segments for each segment pair. If no predicted segment overlaps with the annotated segment, we count it as an “unpaired segment”. The percent of boundaries within 10 amino acids of the annotated segments is relative to the total number of annotations, even if the annotated segment does not have an overlapping predicted segment (unpaired segments are counted towards the total percentage).

### Transferring UniProt annotations to label segment embeddings

We transfer annotations from UniProt to segments defined by ZPS based on how much the segment overlaps with a UniProt annotation. When labeling segments, we allow UniProt annotations of any size, as opposed to evaluating segmentation/boundary prediction only allow annotations that are 30 amino acids or larger. We handle the many overlapping annotations that can be found on UniProt by selecting the annotation that has the highest IoU with the ZPS protein segment. An annotation is only considered to be transferred if the intersection between the UniProt annotation and the ZPS protein segment is at least 30% of the ZPS protein segment.

When we consider annotations that are overlapping IDRs to assess segment embeddings of IDRs, we first define ZPS segments as IDRs if at least 30% of their amino acids overlap with a disordered annotation from MobiDB, then we perform the annotation transfer described above only for ZPS segments that are at least partially IDRs.

### Nearest neighbours

Here we use nearest neighbour to identify similar segment embeddings using cosine distance. We use this to predict the annotations of segment embeddings by finding the label of their 1-nearest neighbour (1-nn) and evaluate with a normalized confusion matrix (see below). We also use nearest neighbours to find protein segments that are similar to the SYGQ-rich prion-like domain in FUS (Fig 7).

### Normalized Confusion Matrix

We use cosine similarity to identify the 1-nn of protein segment embeddings and use annotation of the 1-nn as the predicted label. We do not allow a segment to have the predicted label from another segment from the same protein; we simply choose the next nearest neighbour that is not from the same protein. We use this to define a confusion matrix which is normalized by the number of segments with that label. We also report a binomial proportion confidence interval for values along the diagonal. When we perform 1-nn to evaluate on a specific set of labels, such as the top 20 ProRule domain annotations, only segments with those annotations are included in the evaluation.

### PyMOL

We use open-source Pymol available on github. We used their interactive interface to colour protein structures by protein segments defined by ZPS (Figs 2C, 4D, 4E).

The PyMOL Molecular Graphics System, Version 2.5.0 Schrödinger, LLC. https://github.com/schrodinger/pymol-open-source

### Clustergrams with Plotly

We use clustergrams available through Plotly DashBio application to generate heatmaps of ProtT5 embeddings (Fig 1A) and perform clustering (Fig 1A, 2B). We cluster using hierarchical agglomerative clustering with average linkage and cosine distance; we also use optimal leaf ordering.

### Clustering unannotated segments with Leiden

We use ScanPy [44] to perform Leiden clustering [45] of unannotated ZPS segments from the human proteome. We do not reduce the dimensionality of the segment embeddings before clustering, and we allow for the model to find the optimal clusters; we do not limit the number of iterations it runs and chose a cluster resolution of 2.

### GO enrichment (PANTHER)

To determine which proteins localized to the mitochondria, we performed a GO Enrichment using PantherDB [62] (Version 19.0, released 2024-06-20). We report FDR corrected p-values.

## Acknowledgements

We would like to acknowledge Alex Lu, Mica Consens, Alex Huang, and Andrew Duncan for their valuable feedback on the manuscript.

## Data and Code Availability

The following data used for this analysis is available on Zenodo. https://zenodo.org/records/14962518

1. Human protein sequences that were used to generate per-residue ProtT5 embeddings and segment embeddings.
2. UniProt annotations that were used to evaluate ZPS and label segment embeddings.
3. A list of segment boundaries for the human proteome defined by ZPS.
4. Segment embeddings for the human proteome.

Code to generate per-residue ProtT5 embeddings, define the boundaries of protein segments using change point analysis, and generate segment embeddings is available on GitHub and can be run on colab.

https://github.com/moses-lab/zero-shot-protein-segmentation

Code to visualize protein segment embeddings from the human proteome as colours along the protein’s sequence is available on GitHub and can be run on colab. https://github.com/moses-lab/pLM-Visualization

## Supporting Information

**S1 Table. 1-Nearest Neighbour Normalized Confusion Matrix Accuracy and Binomial Proportion Confidence Intervals.** See separate tables within the excel file for the different 1-nn confusion matrices. This includes tables for IDR vs domain, 5 compositional biases, protein domains, protein sub-domains, and IDRs as well as the corresponding 1-mer (one hot encoding) or 3-mer tables, for instances when they are compared to these in the main text.

**S2 Table. Protein Identifiers and Segment Boundaries of Protein Kinase Domain Segment Clusters.** Identifiers are UniProt IDs, boundaries use zero-based indexing and labels include a, b, c, and d as shown in Fig 4. These segments have not been corrected for over-segmentation, meaning there can be multiple segments from the same proteins (UniProt IDs are non-unique).

**S3 Table. Protein Identifiers and Segment Boundaries of Protein Segments in the Mitochondrion Analysis.** Identifiers are UniProt IDs, boundaries use zero-based indexing and labels include “Annotated” (only includes protein segments that overlap with Mitochondrion Related cluster in the UMAP, Fig 6A) and “10” (corresponding to Leiden cluster number 10, labelled as “Mitochondrion Related” in cyan in Fig 6). These segments have not been corrected for over-segmentation, meaning there can be multiple segments from the same proteins (UniProt IDs are non-unique).

**S4 Table. Protein Identifiers and Segment Boundaries of Protein Segments Similar to FUS’s SYGQ-rich Prion-Like Domain.** Identifiers are UniProt IDs and boundaries use zero-based indexing. These segments have been corrected for over-segmentation, meaning “POS” contains a list of start and stop boundaries of each segment for each protein.

